# Identification of Fibronectin 1 as a candidate genetic modifier in a *Col4a1* mutant mouse model of Gould syndrome

**DOI:** 10.1101/2020.11.06.371666

**Authors:** Mao Mao, Tanav Popli, Marion Jeanne, Kendall Hoff, Saunak Sen, Douglas B. Gould

## Abstract

Collagen type IV alpha 1 and alpha 2 (COL4A1 and COL4A2) are major components of almost all basement membranes. *COL4A1* and *COL4A2* mutations cause a multisystem disorder called Gould syndrome which can affect any organ but typically involves the cerebral vasculature, eyes, kidneys and skeletal muscles. The manifestations of Gould syndrome are highly variable and animal studies suggest that allelic heterogeneity and genetic context contribute to the clinical variability. We previously characterized a mouse model of Gould syndrome caused by a *Col4a1* mutation in which the severities of ocular anterior segment dysgenesis (ASD), myopathy, and intracerebral hemorrhage (ICH) were dependent on genetic background. Here, we performed a genetic modifier screen to provide insight into the mechanisms contributing to Gould syndrome pathogenesis and identified a single locus (*modifier of Gould syndrome 1*; *MoGS1*) on Chromosome 1 that suppressed ASD. A separate screen showed that the same locus ameliorated myopathy. Interestingly, *MoGS1* had no effect on ICH, suggesting that this phenotype may be mechanistically distinct. We refined the *MoGS1* locus to a 4.3 Mb interval containing 18 protein coding genes, including *Fn1* which encodes the extracellular matrix component fibronectin 1. Molecular analysis showed that the *MoGS1* locus increased *Fn1* expression raising the possibility that suppression is achieved through a compensatory extracellular mechanism. Furthermore, we show evidence of increased integrin linked kinase levels and focal adhesion kinase phosphorylation in *Col4a1* mutant mice that is partially restored by the *MoGS1* locus implicating the involvement of integrin signaling. Taken together, our results suggest that tissue-specific mechanistic heterogeneity contributes to the variable expressivity of Gould syndrome and that perturbations in integrin signaling may play a role in ocular and muscular manifestations.

## INTRODUCTION

Collagens are the most abundant proteins in the body, making up ~30% of the dry weight. The collagen superfamily of extracellular matrix (ECM) molecules comprises 28 members encoded by 46 genes (Ricard-Blum, 2011). Among these, type IV collagens are primordial ECM molecules and are fundamental constituents of specialized structures called basement membranes (Fidler et al., 2017). In mammals, six genes encode the type IV collagens (*COL4A1* to *COL4A6*) and their protein products assemble into three distinct heterotrimers (α1α1α2, α3α4α5 or α5α5α6). The α1α1α2 network is ubiquitous throughout development and in most adult tissues and its absence results in embryonic lethality (Poschl et al., 2004). Pathogenic mammalian *Col4a1* mutations were first identified using forward mutagenesis screens in mice with variable forms of ocular pathology (Favor et al., 2007; Gould et al., 2005; Thaung et al., 2002) and human mutations were identified in individuals with severe inherited or *de novo* porencephaly and early onset intracerebral hemorrhages (ICH) (Breedveld et al., 2006; Gould et al., 2005; Gould et al., 2006; Sibon et al., 2007; Vahedi et al., 2007). A number of studies in humans and mice have subsequently expanded the phenotypic spectrum and it is now well established that mutations in *COL4A1* and *COL4A2* cause a multi-system disorder (Jeanne and Gould, 2017; Labelle-Dumais et al., 2019; Mao et al., 2015; Meuwissen et al., 2015; Yoneda et al., 2013; Zagaglia et al., 2018) that has adopted the name Gould syndrome (https://www.gouldsyndromefoundation.org).

Gould syndrome is highly clinically heterogeneous and can have variable penetrance and severity across many organs. Cerebrovascular, ocular, renal and neuromuscular pathologies are among the most commonly described manifestations. We originally described a mouse model of Gould syndrome with a *Col4a1* splice acceptor mutation that leads to exclusion of exon 41 (∆ex41) from the mature transcript and 17 amino acids from the triple-helical domain of the protein (Gould et al., 2005). However, the majority of disease-causing variants reported in humans are missense mutations in highly conserved glycine residues of the triple-helical domain (Jeanne and Gould, 2017). In order to faithfully replicate human disease, we compared the effects of different mutations in an allelic series composed of nine distinct *Col4a1* and *Col4a2* mutant mouse strains (Favor et al., 2007; Jeanne et al., 2015; Kuo et al., 2014). We established that allelic heterogeneity has important implications for penetrance and severity of various pathologies including ICH and myopathy. We found that mutations can differ in degree or kind and that allelic differences contribute to clinical variability, in part, by tissue-specific mechanistic heterogeneity (Labelle-Dumais et al., 2019). However, variable clinical manifestations including age of onset and severity among organs have been reported in people with the same recurrent or inherited mutations, and reduced penetrance and asymptomatic carriers exist in some families, indicating the involvement of factors in addition to allelic heterogeneity (Coupry et al., 2010; de Vries et al., 2009; Rødahl et al., 2013; Shah et al., 2012).

Studies using inbred strains of mice clearly demonstrate that a mutation can have variable outcomes in different genetic contexts underscoring the important role that genetic modification plays in clinical heterogeneity. When compared to other mutations in the allelic series, *Col4a1*^*Δex41*^ tends to have the most severe phenotypes across multiple organs when maintained on a pure C57BL/6J (B6) genetic background (Gould et al., 2007; Jeanne et al., 2015; Kuo et al., 2014). However, when the *Col4a1*^*+/Δex41*^ B6 mice were crossed for a single generation to the CAST/EiJ (CAST) genetic background, ocular anterior segment dysgenesis (ASD), ICH and skeletal myopathy were all significantly reduced in the F1 progeny (CASTB6F1) (Gould et al., 2007; Jeanne et al., 2015; Labelle-Dumais et al., 2011). Interestingly, when *Col4a1*^*+/Δex41*^ B6 mice were crossed for a single generation to the 129SvEvTac (129) genetic background, ASD was ameliorated (albeit to a lesser extent than in CASTB6F1 mice) but ICH was not (Gould et al., 2007; Jeanne et al., 2015). Collectively, these data suggest that the B6 genetic background confers susceptibility to develop severe Gould syndrome related pathologies and that CAST and 129 strains have one or more locus/loci that can genetically suppress pathology in a tissue-specific manner.

Identification of genetic modifier genes can help reveal disease mechanisms and potential therapeutic targets. Phenotype-driven mapping studies using mice and other model organisms are a powerful approach to identify genetic interactions that may be difficult to discover in humans even when large pedigrees are available (Ceco and McNally, 2013; Meyer and Anderson, 2017; Vieira et al., 2015). We previously performed a genetic modifier screen for pathology caused by the *Col4a1*^*Δex41*^ mutation to provide insight into the underlying pathogenic mechanisms. Because 1) ASD is relatively severe, easily screened, and robustly suppressed by two strains and 2) the magnitude and breadth of phenotypic rescue was greater in CAST compared to 129 mice, we selected ASD and CAST as the phenotype and strain, respectively, to perform a pilot suppressor screen in a small number of mice (Gould et al., 2007). We generated CASTB6F1 mice and iteratively crossed mutant mice back to the B6 background. In each backcross, we applied selective pressure to retain one or more genetic suppresser locus by choosing the progeny with the mildest phenotype to breed for the next generation. Using a crude genome-wide scan, we identified a dominant modifier locus in CAST Chromosome (Chr) 1 that suppressed ASD. However, the approach had significant limitations including screening on a single phenotype, the inability to identify potential recessive loci and the possibility that we may have incidentally overlooked other dominant loci by imposing bottlenecks at each generation.

Here, we performed independent genetic modifier screens for ASD and skeletal myopathy on a large scale F2 cross with the potential to identify multiple dominant or recessive loci that enhanced or suppressed pathology. Both screens identified a single suppressor locus on CAST Chr 1 that we called *modifier of Gould syndrome 1* (*MoGS1*). Surprisingly, although the CAST background also suppressed ICH, the effect was not attributable to the *MoGS1* locus. These data suggest that ASD and skeletal myopathy may be mechanistically related to each other but distinct from ICH which further supports the notion of tissue-specific mechanistic heterogeneity contributing to the clinical variability of Gould syndrome (Labelle-Dumais et al., 2019). Furthermore, molecular analyses suggest that the ECM protein fibronectin 1 (FN1) is a strong candidate as the genetic suppressor at the *MoGS1* locus and provide evidence for a role of altered integrin signaling in Gould syndrome pathogenesis.

## RESULTS

### High-resolution genome-wide mapping identified a suppressor locus for *Col4a1*-related ASD and myopathy on mouse Chr 1

To identify modifiers of Gould syndrome, we first performed a genome-wide screen with the early onset and easily observed ASD phenotype. We phenotyped and genotyped 192 mutant F_2_ progeny from (CAST X B6-*Col4a1*^*+/Δex41*^) F_1_ intercrosses using a mouse medium density linkage panel with 646 informative SNPs (Fig. 1A-C). We identified a region of interest on CAST Chr 1 with a LOD score of 11.2 and Bayesian confidence interval extending from 51.3 to 73.0 Mb (Fig.1C). To determine if smaller effects of other loci might have been masked, we performed a second genome-wide scan conditioned on this locus, but no additional loci reached statistical significance (Fig. 1D). This observation suggests that the major modifying effect of the CAST background on ASD is imparted through this single dominant locus on Chr 1.

**Figure 1.**
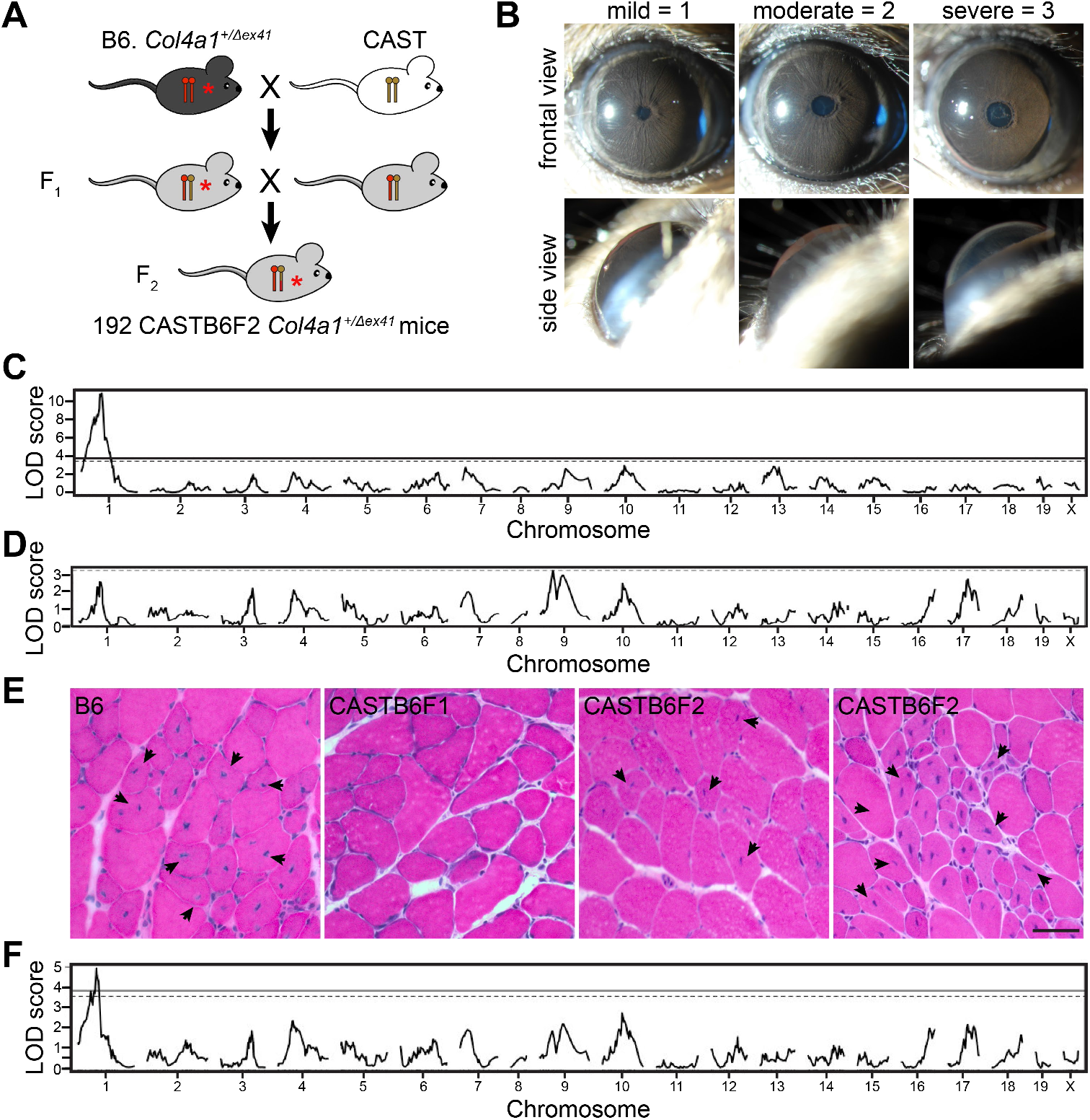
A genome-wide screen identified a single modifier locus on Chr 1 for ASD and myopathy. **(A**) Schematic representation of the (CAST X B6) F2 cross. One hundred ninety two mutant (CAST X B6-*Col4a1*^*+/Δex41*^) F_2_ progeny were genotyped and phenotyped for ASD. (**B**) Representative slit-lamp images illustrating scoring of ASD severity: mild, score = 1; moderate, score = 2; severe, score = 3. (**C**) A one-dimensional genome scan identified a locus on Chr 1 for ASD (max. LOD score = 11.2, 99% Bayesian confidence interval extending from 51.3 - 73.0 Mb, Ensembl GRCm38.p6). Solid horizontal line, 5% false positive threshold; dashed horizontal line, 10% threshold. (**D**) A genome-wide scan conditioned on the Chr 1 modifier locus suggests that no other single locus has a strong effect on ASD. Dashed horizontal line: 20% false positive threshold. (**E**) Representative images of cross sections from quadriceps stained with H&E showing variable myopathy severity in *Col4a1*^*+/Δex41*^ mice with different genetic backgrounds. Arrows indicate muscle fibers with NPN. Scale bar = 20 μm. (**F**) To identify skeletal muscle modifier loci, a subset (49) of the 192 (CAST X B6) F2 progeny was assessed for myopathy. One dimensional genome scan identified a myopathy modifier at the same position as the ASD modifier (Max. LOD score = 5.61).

In addition to suppressing ASD, the CAST genetic background significantly suppresses skeletal myopathy in *Col4a1* mutant mice (Labelle-Dumais et al., 2011). To identify potential modifier loci for this phenotype, we performed an independent genome-wide mapping analysis using skeletal myopathy as the primary phenotype. We quantified myopathy severity as the percentage of muscle fibers containing non-peripheral nuclei (NPN) in histological sections of quadriceps (Fig. 1E). This assay is more quantitative with greater statistical power but also more labor-intensive and therefore we analyzed a randomly generated subset (n= 49) of the mutant (CAST X B6-*Col4a1*^*+/Δex41*^) F2 progeny. The genome-wide scan revealed an interval on Chr 1 defined by the same markers as the ASD modifier locus (max LOD score = 5.61) (Fig. 1F).

To validate the biological effect of the *MoGS1* locus, we independently generated an incipient congenic strain by iterative backcrossing the CAST-derived locus onto the susceptible B6 genetic background for 5 generations (N5). At N5, the genetic background of the mice is ~97% pure B6 on average, except that they carry the CAST Chr 1 interval that includes the *MoGS1* locus. Incipient congenic mice were intercrossed to generate *Col4a1*^*+/Δex41*^ mice with zero (B/B), one (C/B) or two (C/C) copies of the CAST-derived chromosomal interval (Fig. 2). Out of 24 eyes from *Col4a1*^*+/Δex41*^ mice that were homozygous for the B6 allele (*MoGS1*^*B/B*^) at the congenic interval, 29% (7) were moderate, and 71% (17) were severe (Fig. 2A). In contrast, out of 28 eyes from the heterozygous group (*MoGS1*^*C/B*^) there were 21.5% (6), 46.5% (13) and 32% (9) of eyes that were scored with mild, moderate and severe ASD, respectively. In mice homozygous for the CAST allele (*MoGS1*^*C/C*^), there were 50% (10), 35% (7) and 15% (3) of eyes that were mild, moderate and severe, respectively. In a parallel experiment, we tested the effect of this congenic locus on myopathy and observed a dosage effect, whereby *Col4a1*^*+/Δex41*^ mice that were heterozygous at the *MoGS1* locus (*MoGS1*^*C/B*^) showed a trend toward reduced myopathy while homozygous *MoGS1*^*C/C*^ incipient congenic mice had significantly milder myopathy compared to mice that were homozygous for the B6 allele (*MoGS1*^*B/B*^). Together, these results support the existence of one or more semi-dominant modifier genes at the *MoGS1* locus on Chr 1.

**Figure 2.**
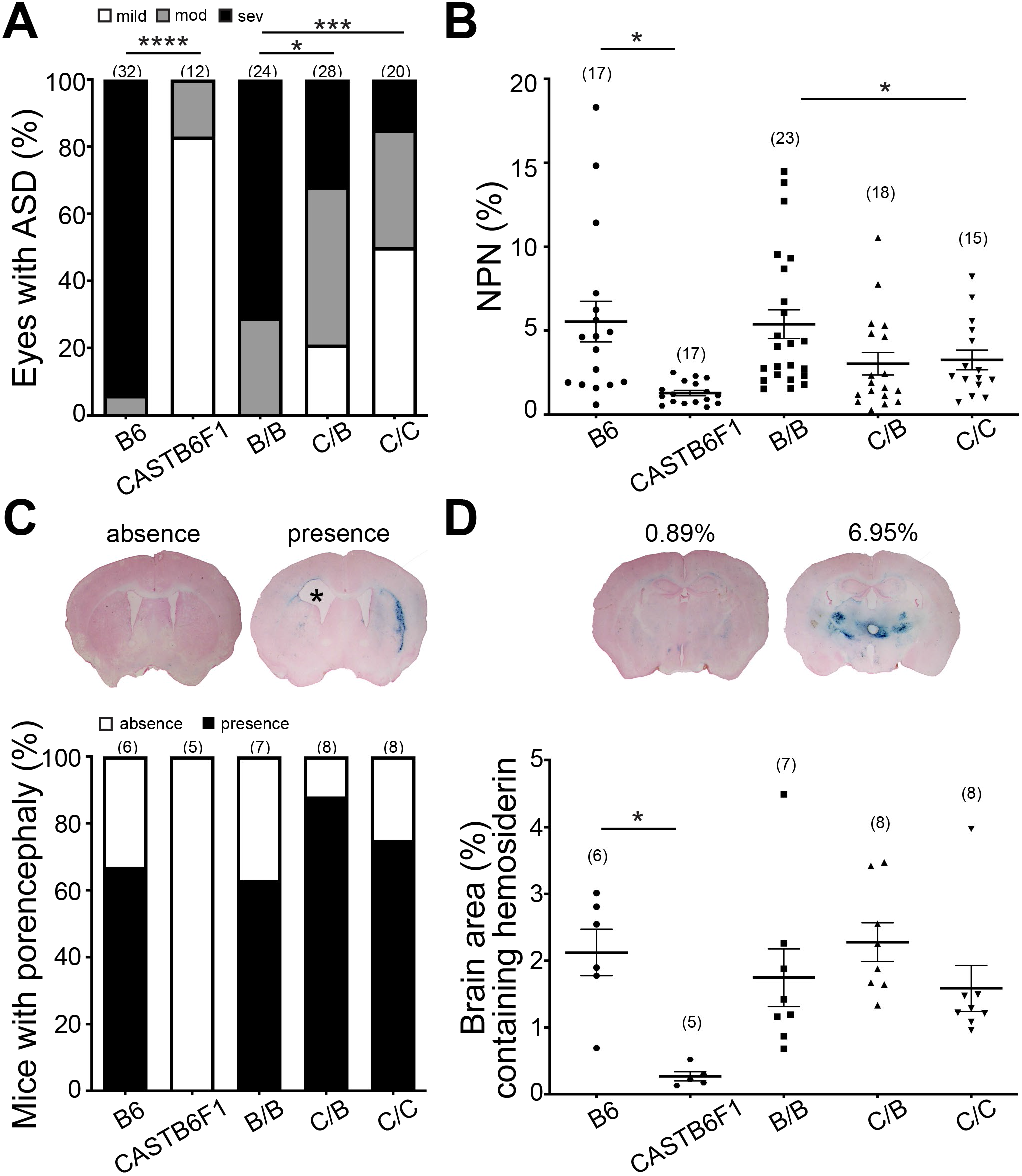
An incipient congenic strain containing the CAST-derived *MoGS1* locus suppressed ASD and myopathy, but not ICH. (**A**) Evaluation of eyes from the incipient congenic mice at N5F2 validates *MoGS1* as a genetic suppressor of ASD. The bar graph shows the percentage of eyes at each ASD severity level from *Col4a1*^*+/Δex41*^ mice that are homozygous for B6 alleles (B/B), heterozygous or homozygous for CAST alleles (C/B and C/C, respectively) in the congenic interval. ASD for *Col4a1*^*+/Δex41*^ mice on the pure B6 and CASTB6F1 backgrounds are also shown. \**p* < 0.05; \*\*\*\**p* < 0.0001, by Kruskal-Wallis test followed by Dunn’s multiple comparison test. (**B**) Evaluation of muscles from the N5F2 congenic mice validates *MoGS1* as a genetic suppressor of skeletal myopathy. Data are presented as mean ± SEM. \**p* < 0.05, by one-way ANOVA followed by Tukey’s multiple comparison test. (**C**) Penetrance of porencephaly for *Col4a1*^*+/Δex41*^ mice with different genetic backgrounds was determined by brain histology. Top representative images show absence or presence of porencephaly. * marks brain area with porencephaly. (**D**) ICH in mice was assessed using Perl’s Prussian blue staining. Data for each mouse are shown as a percentage of brain area with Prussian blue staining averaged over 28 sections from each brain. Representative images of Prussian blue-stained brain sections and respective Prussian blue staining quantification are shown above the graphs. Sample sizes are indicated in parentheses. Data are presented as mean ± SEM. \**p* < 0.05, Student’s t-test for comparison between B6 and CASTB6F1; One-way ANOVA followed by Tukey’s test for multiple comparisons between B/B, C/B, and C/C.

### *MoGS1* does not reduce porencephaly penetrance or ICH severity in *Col4a1* mutant mice

Individuals with Gould syndrome have highly penetrant and clinically variable cerebrovascular diseases that include porencephaly and ICH (Bilguvar et al., 2009; de Vries et al., 2009; Giorgio et al., 2015; Vilain et al., 2002). ICH was previously reported to be significantly reduced in *Col4a1* mutant mice on a CASTB6F1 background compared to those maintained on a B6 background suggesting that the CAST genetic background also suppresses this phenotype (Jeanne et al., 2015). To test whether the *MoGS1* locus might also genetically modify cerebrovascular diseases, we assessed porencephaly penetrance and ICH severity of the incipient congenic mice. We found that porencephaly penetrance and ICH severity were similar among *Col4a1*^*+/Δex41*^ mice irrespective of their genotypes at the *MoGS1* locus (Fig. 2C, D).

### Fine mapping with subcongenic lines refined *MoGS1* to a 4.3 Mb interval

To refine the interval of interest at the *MoGS1* locus, we generated subcongenic lines of the CAST derived chromosomal fragments on a B6 background (N5) and tested which line suppresses ASD and skeletal myopathy in *Col4a1*^*+/Δex41*^ mice (Fig. 3A). Line 1 which included an interval of ~10 Mb from the CAST genome did not have a modifying effect on either phenotype (Fig. S1). In contrast, a subcongenic line (line 2) containing the distal portion of the *MoGS1* locus (from 68.7 – 73.0 Mb) showed a significant protective effect for both ASD (Fig. 3B) and myopathy (Fig. 3C). While approximately 87% of eyes from *Col4a1*^*+/Δex41*^;*MoGS1*^*B/B*^ mice had severe ASD (1 mild, 4 moderate and 33 severe), only ~35% of the eyes from *Col4a1*^*+/Δex41*^;*MoGS1*^C/C^ mice were severe (8 mild, 14 moderate, and 12 severe). Likewise, *Col4a1*^*+/Δex41*^;*MoGS1*^*C/C*^ mice had significantly fewer NPN compared to *Col4a1*^*+/Δex41*^;*MoGS1*^*B/B*^ mice. Heterozygosity for this distal interval had intermediate effects for both phenotypes.

**Figure 3.**
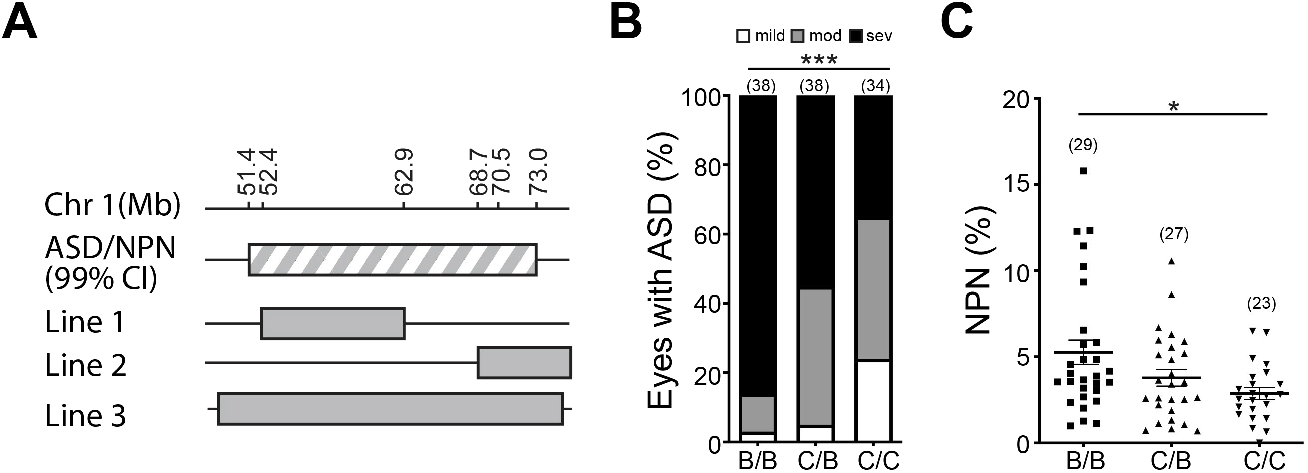
Fine mapping of the *MoGS1* locus. (**A**) Schematic illustration of subcongenic lines containing different intervals of the CAST modifier locus backcrossed onto the B6 background for 5 generations. Subcongenic Line 2 (68723089 – 72973981bp on Chr 1, Ensembl GRCm38.p6) ameliorated ASD and myopathy severity. Black line, chromosome fragments from B6; grey bar, chromosome fragments from CAST. CI, critical interval. (**B**) Analyses of ASD severity in mice homozygous for B6 alleles (B/B), heterozygous or homozygous for CAST alleles (C/B and C/C, respectively) at the refined distal interval. Percentage of eyes at each ASD severity level was shown. \*\*\**p*<0.001 by Kruskal-Wallis test followed by Dunn’s multiple comparison test. (**C**) Analysis of myopathy by NPN showing severity in mice with different genotypes at the refined distal interval. Sample size indicated in parentheses. Data are presented as mean ± SEM.\**p* < 0.05, one-way ANOVA followed by Tukey’s multiple comparison test.

### Candidate gene analysis

The refined *MoGS1* locus is 4.3 Mb, and contains 18 protein-coding genes, and 38 predicted non-protein coding genes including 13 lncRNAs, 6 snRNA genes, pseudogenes, and others (Table S1). Using publicly available databases (Keane et al., 2011), we examined the refined *MoGS1* interval for sequence differences between the B6 and CAST genomes. Among all variations, there were 7,430 single nucleotide polymorphisms (SNPs), 667 small insertions and deletions (indels) and 17 large structural variations including large insertions or deletions. Thirty-five SNPs were predicted to be missense variants affecting 10 protein-coding genes (Table 1). Among those genes, *Ankar* had three missense variants predicted to be deleterious by SIFT (Vaser et al., 2016) and *Spag16* had one predicted stop-gain variant; however, neither gene is an obvious functional candidate. Based on the Gene Expression Database (GXD) from Mouse Genome Informatics (informatics.jax.org/expression), 13 of the 18 protein coding genes have been shown to be expressed both in eyes and skeletal muscles. Two genes, *Vwc2l* and *Bard1*, are only expressed in eye and three genes, *Ankar*, *Spag16*, and *Abca12*, have no reported expression in either tissue. Since both ocular and muscular defects in *Col4a1*^*+/Δex41*^ mice were suppressed by the *MoGS1* locus, we hypothesized that the underlying pathogenic mechanism(s) may be shared between these two tissues. Therefore, despite the presence of possible pathogenic variants, we excluded *Ankar, Spag16*, *Abca12, Vwc2I and Bard1* from further analysis.

**Table 1.**
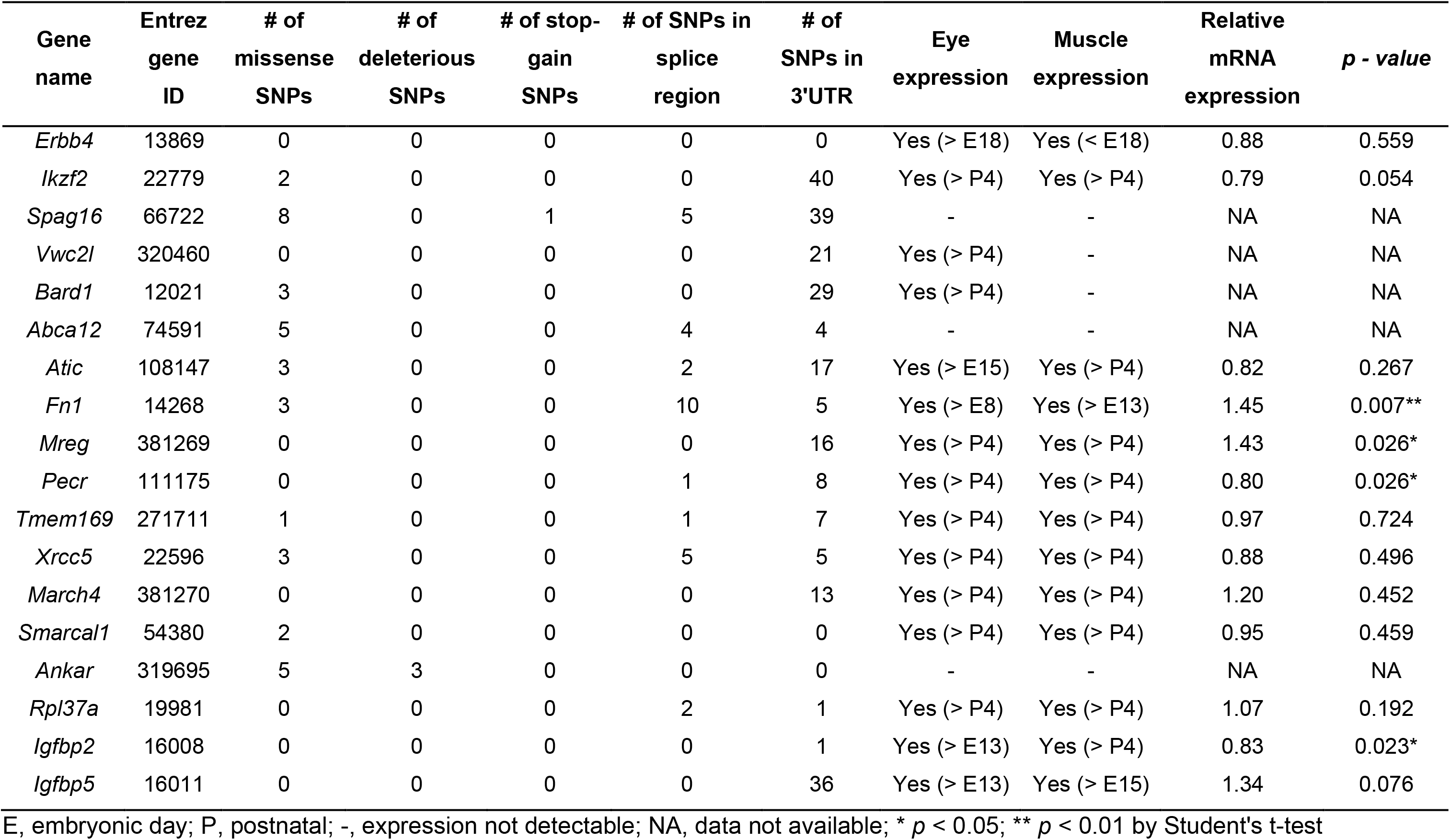
List of protein-coding genes in the refined *MoGS* locus.

Next, we tested whether any of the 13 genes with reported expression in ocular and muscular tissues are differentially expressed in B6 mice with or without the CAST interval at the *MoGS1* locus. Four genes, *Mreg*, *Fn1*, *Pecr1*, and *Igbp2*, showed significantly increased expression in *MoGS1^C/C^* eyes at postnatal day 0 (P0) compared to *MoGS1*^*B/B*^ eyes by qPCR (Table 1 and Fig. 4A). FN1 is an extracellular matrix protein with multiple important roles in development and tissue homeostasis by interacting with cell surface receptors, extracellular matrix proteins and growth factors (Zollinger and Smith, 2017). Notably, FN1 has binding sites for type IV collagen (Laurie et al., 1986; Laurie et al., 1982) and localizes adjacent to basement membranes in many tissues (Laurie et al., 1982). Moreover, like collagens, FN1 interacts with cells via integrin receptors (Johansson et al., 1997; Vandenberg et al., 1991), making it a strong functional candidate as a genetic modifier of COL4A1-related pathology.

**Figure 4.**
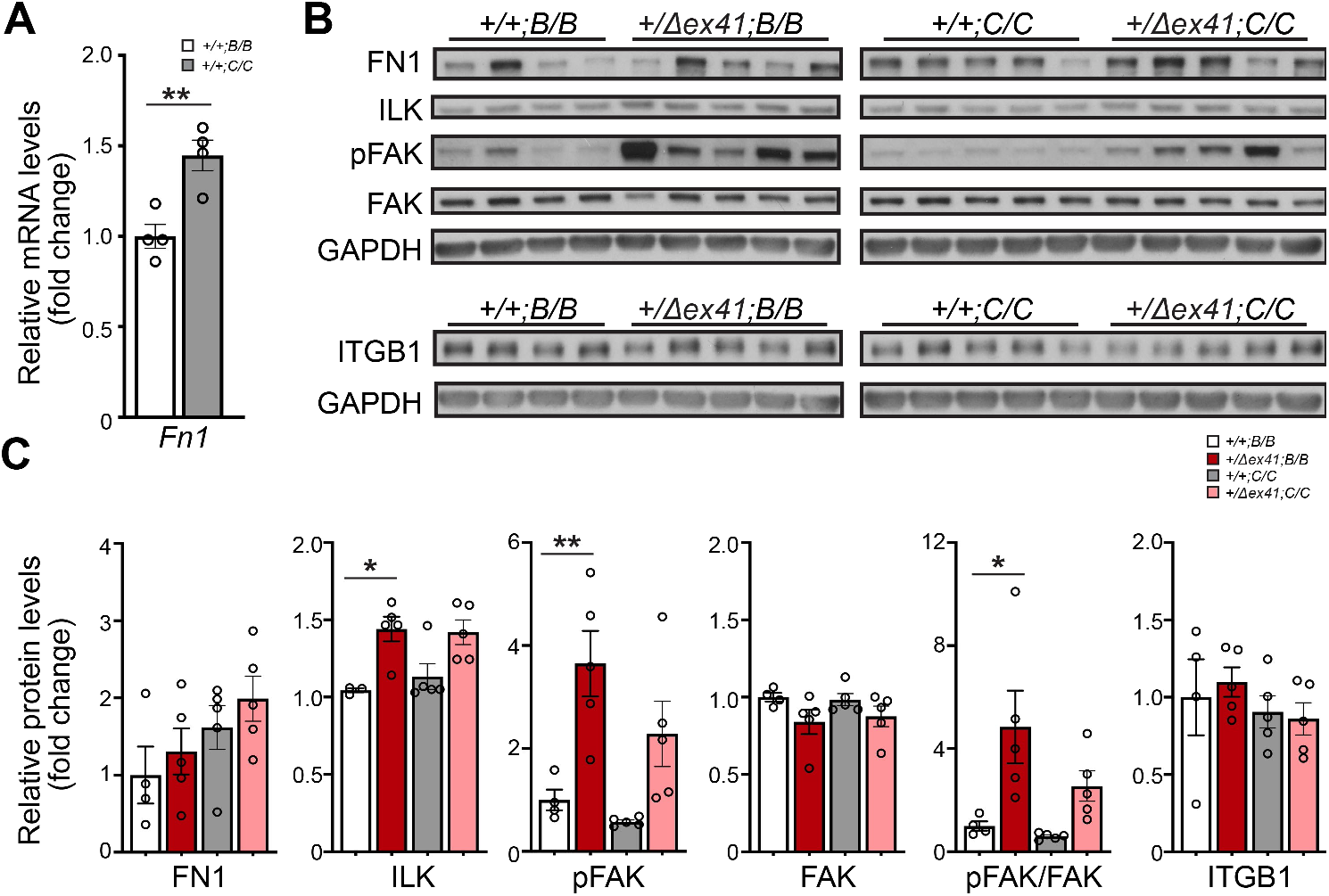
Evaluating FN1 as a candidate modifier gene. (**A**) Quantitative qPCR analysis showed increased expression of *Fn1* mRNA in P0 eyes from *Col4a1*^*+/+*^ mice homozygous for the CAST interval at the refined *MoGS1* locus (*+/+;C/C*) compared to mice homozygous for the B6 interval (*+/+;B/B*). *n* = 4 per genotype. Data are presented as mean ± SEM. \**p* < 0.05, by Student’s t-test. (**B-C**) Representative images (B) and quantification (C) of Western-blot analyses of P10 quadriceps showing a trend towards increased FN1 protein levels in mice homozygous for CAST alleles at the refined *MoGS1* locus *(+/+;C/C* and *+/Δex41;C/C*) compared to mice homozygous for B6 alleles (*+/+;B/B* and *+/Δex41;B/B*). ILK levels were increased in *Col4a1^+/Δex41^* mice irrespective of the genotype at the *MoGS1* locus (*+/Δex41;B/B* and *+/Δex41;C/C*, *p = 0.06* for *MoGS1*^*C/C*^) compared to their corresponding *Col4a1*^*+/+*^ controls (*+/+;B/B, and +/+;C/C,* respectively). Ratios of pFAK/FAK, were significantly elevated in *Col4a1*^*+/Δex41*^ mice homozygous for *MoGS1* B6 alleles *(+/Δex41;B/B*) but not CAST alleles (*+/Δex41;C/C*) (*p* = 0.35) compared to their corresponding *Col4a1*^*+/+*^ controls (*+/+;B/B, and +/+;C/C,* respectively) suggesting altered integrin signaling that is partially restored by the *MoGS1* locus. Levels of ITBG1 did not differ between genotypes. *n* = 4, 5, 5, 5 for *+/+;B/B*, *+/Δex41;B/B*, *+/+;C/C* and *+/Δex41;C/C* respectively. Data are presented as mean ± SEM. \**p* < 0.05, \*\**p* < 0.01 by one-way ANOVA followed by Sidak’s multiple comparison test.

### Functional testing of FN1 and integrin signaling in MoGS1 mice

In silico analysis comparing the CAST and B6 alleles of *Fn1* revealed 3 missense SNPs, 1 splice region SNP, 7 SNPs in the UTR, 238 other noncoding SNPs, and 2 structural variations (Table S2). Murine FN1 has 12 isoforms (White et al., 2008), making it difficult to predict the functional consequence(s) of a particular variant. The primary consequence of *Col4a1* mutations is impaired secretion of mutant α1α1α2 heterotrimers into the basement membranes (Kuo et al., 2014) and it is possible that increased FN1 confers partial compensation for extracellular α1α1α2 deficiency. Consistent with this hypothesis, we observed increased *Fn1* expression in developing eyes from *MoGS1*^*C/C*^ mice compared to *MoGS1*^*B/B*^ mice (Fig. 4A). Furthermore, we found that FN1 levels were higher in P10 quadriceps from *Col4a1*^*+/Δex41*^ compared to *Col4a1*^*+/+*^ mice and in *MoGS1*^*C/C*^ compared to *MoGS1*^*B/B*^ mice (Fig. 4B, C). FN1 plays important roles in cell adhesion, migration and signaling during tissue morphogenesis, which is primarily mediated through its interaction with integrins (Miyamoto et al., 1998; Sakai et al., 2003). Therefore, we tested the effect of *MoGS1* on integrin signaling by evaluating the protein levels and phosphorylation status of two downstream effectors, integrin linked kinase (ILK) and focal adhesion kinase (FAK) (Hu and Luo, 2013; Humphries et al., 2019) (Fig. 4B, C). Western blot analysis of P10 quadriceps revealed that ILK levels were higher in *Col4a1*^*+/Δex41*^ mice irrespective of whether they carried the *MoGS1*^*B/B*^ or *MoGS1*^*C/C*^ interval. Similarly, FAK phosphorylation (pFAK) and pFAK/FAK ratio were significantly higher in *Col4a1*^*+/Δex41*^;*MoGS1*^*B/B*^ compared to *Col4a1*^*+/+*^;*MoGS1*^*B/B*^ mice, and were reduced by the *MoGS1*^*C/C*^ interval. No difference was detected for the broadly used beta subunit of the integrin receptor, ITGB1, between genotypes. Taken together, these data suggest that elevated integrin signaling may contribute to Gould syndrome and that ASD and myopathy may be partially rescued by compensatory *Fn1* expression.

## DISCUSSION

Here, we performed a genetic modifier screen in *Col4a1* mutant mice to gain insight into the pathogenic mechanisms that contribute to Gould syndrome. To this end, we used a large-scale F2 cross of CASTxB6 and conducted independent screens for ASD and skeletal myopathy – two phenotypes that are commonly observed in individuals with Gould syndrome. Independent analyses for both phenotypes revealed a single, shared, semi-dominant locus on CAST Chr 1 that corroborates a previously identified locus (Gould et al., 2007). To validate this genetic data, we iteratively backcrossed the locus onto the B6 background and showed that a refined interval, referred to as *MoGS1*, suppressed ASD and myopathy in *Col4a1* mutant mice. However, the *MoGS1* locus had no effect on porencephaly penetrance and ICH severity – two phenotypes associated with Gould syndrome and previously characterized in *Col4a1* mutant mice. The failure of *MoGS1* to suppress ICH severity was unexpected since the CASTB6F1 genetic context significantly suppresses this phenotype. These data suggest that the effect of the *MoGS1* locus is tissue-specific and that other modifiers of ICH exist in the CAST background.

Using subcongenic strains we refined the minimum critical interval for the suppressor locus to a 4.3 Mb region containing 18 protein-coding genes. Of the 18 positional candidate genes, 13 are expressed both in eyes and muscles, of which 6 contained missense variants between the CAST and B6 genomes; however, none were predicted to be deleterious. Mutations in one of these six genes, *Smarcal1*, causes a rare multisystem disorder (Schimke Immunoosseous dysplasia; OMIM# 242900) characterized by spondyloepiphyseal dysplasia, nephrotic syndrome, T cell immunodeficiency, and a portion of patients develop corneal opacity, myopia, astigmatism and optic atrophy (Boerkoel et al., 2000). SMARCAL1 (SWI/SNF related matrix associated, actin dependent regulator of chromatin, subfamily a-like 1) is chromatin remodeling protein involved in transcriptional regulation and DNA replication, repair and recombination (Bansbach et al., 2010). Compared to the B6 reference genome, CAST has two missense variations in *Smarcal1* that were predicted to be tolerated, and no change of expression was detected in P0 eyes. Of the four positional candidate genes that were differentially expressed, *Fn1* is a strong functional candidate. FN1 is a ubiquitously expressed extracellular glycoprotein which plays important roles in multiple processes by interacting with cell surface receptors, growth factors and other extracellular matrix proteins. In cell culture, FN1 promotes collagen IV deposition, assembly and incorporation into the extracellular matrix (Filla et al., 2017; Ngandu Mpoyi et al., 2020). Mice deficient for *Fn1* have aberrant lens placode formation and develop microphthalmia and cataracts (Hayes et al., 2012; Huang et al., 2011). Moreover, lack of FN1 in the skeletal muscle stem cell niche impairs muscle regeneration in aged mice (Lukjanenko et al., 2016). Collectively, these findings support our genetic and molecular evidence suggesting that that FN1 is a strong candidate as the gene responsible for the effects conferred by the *MoGS1* locus.

In general, the presence of mutant COL4A1 or COL4A2 leads to intracellular accumulation of mutant heterotrimers at the expense of their secretion. Intracellular heterotrimer accumulation represents a potential cell autonomous (proximal) insult (Jeanne and Gould, 2017; Mao et al., 2015). Impaired heterotrimer secretion can also lead to a cell non-autonomous (distal) insult caused by extracellular heterotrimer deficiency which can simultaneously perturb any number of presently unidentified cellular pathways. Intracellular accumulation and extracellular deficiency can be addressed simultaneously by targeting the proximal defect of protein misfolding. For example, pharmacologically promoting heterotrimer secretion using a chemical chaperone, 4-phenylbutyrate (4PBA), alleviates diverse pathologies in *Col4a1* mutant mice (Hayashi et al., 2018; Jeanne et al., 2015; Jones et al., 2019; Labelle-Dumais et al., 2019). However, even in the absence of 4PBA, secretion of mutant heterotrimers is not completely abolished and the presence of mutant heterotrimers in the basement membrane represents a third and distinct class of insult. We demonstrated this proof of concept by showing that mice with a *Col4a1^G394V^* mutation have disproportionately severe myopathy, which is exacerbated when they are treated with 4PBA (Labelle-Dumais et al., 2019). The affected residue, glycine 394, is adjacent to a putative integrin-binding domain (Parkin et al., 2011) implicating impaired integrin binding in skeletal myopathy. Thus, for mutations that impair subdomains of the heterotrimer responsible for executing specific extracellular functions, promoting secretion of mutant heterotrimers would still be predicted to ameliorate many extracellular functions. However, the increased levels of mutant heterotrimers in basement membranes may also exacerbate specific disease pathway(s) related to the function(s) of the impacted subdomain (Labelle-Dumais et al., 2019). Identifying the multitude of extracellular roles of COL4A1/A2 and how they are executed will be key in understanding fundamental aspects of matrix biology and the tissue-specific pathogenic mechanisms in Gould syndrome. Mapping functional subdomains on the α1α1α2 heterotrimers will be important for genetically stratifying patients and may influence the prognosis and potential therapeutic approaches.

The unbiased nature of genetic screens leaves open the possibility for finding modifiers of either proximal or distal insults. By independently analyzing two phenotypes using an inbred strain (CAST) with broad phenotypic suppression, we sought to find modifiers that might represent therapeutic targets for the breadth of Gould Syndrome phenotypes. *MoGS1*, the only significant locus that we identified, conferred tissue-specific effects by suppressing ASD and myopathy but not ICH. This observation is consistent with a modifier that has an extracellular function and suggests that the two phenotypes have at least partially overlapping distal pathogenic mechanisms. In a previous study using the *Col4a1*^*Δex41*^ mutation we found that, compared to B6, the CASTB6F1 background appeared to reduce intracellular accumulation but did not increase extracellular COL4A1 levels (Jeanne et al., 2015). This observation implicated intracellular accumulation in pathogenesis but did not rule out the potential of compensation by other extracellular factors. The fact that *MoGS1* does not suppress ICH severity indicates that intracellular accumulation of mutant proteins may indeed be the primary pathogenic insult for that phenotype.

When we compared disease severity across a murine allelic series of 9 *Col4a1* and *Col4a2* mutations, we found that the *Col4a1*^*G394V*^ mutation had relatively little intracellular accumulation and that *Col4a1*^*+/G394V*^ mice had mild ICH but severe ASD and myopathy (Kuo et al., 2014). The proximity of the *Col4a1*^*G394V*^ mutation to a predicted integrin binding domain suggests that impaired integrin binding contributes to ocular dysgenesis and myopathy. Therefore, one possibility is that FN1 ameliorates ASD and skeletal myopathy caused by *Col4a1* mutations through compensatory integrin-mediated interactions. Consistent with this hypothesis, we detected evidence for increased integrin signaling in *Col4a1*^*+/Δex41*^ mice that was reduced by the *MoGS1* locus from CAST in the context of increased *Fn1* expression. Moreover, ILK and FAK are both previously implicated in lens development (Cammas et al., 2012; Harburger and Calderwood, 2009; Hayes et al., 2012; Samuelsson et al., 2007; Teo et al., 2014) and myopathy (Boppart and Mahmassani, 2019; Gheyara et al., 2007). Functional validation of this pathway with experimental manipulation and preclinical interventions would represent a significant advance in the understanding of the pathogenic mechanisms that contribute to Gould syndrome.

## MATERIALS AND METHODS

### Mice

All experiments were compliant with the ARVO Statement for the Use of Animals in Ophthalmic and Vision Research and approved by the Institutional Animal Care and Use Committee at the University of California, San Francisco. *Col4a1*^*+/Δex41*^ mice were originally identified in a mutagenesis screen conducted at The Jackson Laboratory (Bar Harbor, ME) (Gould et al., 2007; Gould et al., 2005). Since then *Col4a1*^*+/Δex41*^ mice has been backcrossed to B6 for at least 20 generations. CAST mice were obtained from The Jackson Laboratory. All animals were maintained in full-barrier facilities free of specific pathogens on a 12-hour light/dark cycle with food and water ad libitum. Both male and female mice were used in this study.

### Slit lamp biomicroscopy

Ocular anterior segment examinations were performed on mice at 1.0-1.5 months of age using a slit lamp biomicroscope (Topcon SL-D7; Topcon Medical Systems, Oakland, NJ) attached to a digital SLR camera (Nikon D200; Nikon, Melville, NY).

### Muscle analysis

At 2 months of age, mice were subjected to treadmill exercise and their quadriceps were harvested two days later. Exercise was 30 minutes with a 15° downhill grade on a treadmill equipped with a shock plate (Columbus Instruments, Columbus, OH). Animals were started at 6 m/min and increased by 3 m/min every 2 min until maximum of 15 m/min speed was reached. Quadriceps were dissected and frozen in liquid nitrogen cooled isopentane. Cryosections (10 μm) were collected at regular intervals and stained with hematoxylin and eosin (H&E) for histopathology and the number of NPN were evaluated on a total of 12 sections for each muscle and the percentage of muscle fibers with NPN was quantified. The observers were masked to genotypes and counted between 2000–5000 muscle fibers per animal.

### Brain histological analysis

ICH was assessed at 2 months of age by Perl’s Prussian blue staining as previously described (Jeanne et al., 2015). Briefly, mice underwent transcardial perfusion with saline followed by 4% paraformaldehyde (PFA) and then posted fixed in 4% PFA overnight. Brains were dissected, cryoprotected in 30% sucrose and embedded in optimal cutting temperature compound (Sakura Finetek, Torrance, CA). Coronal cryo-sections (35 μm) regularly spaced along the rostro-caudal axis were stained with Prussian blue/Fast red and imaged. On each section, the percentage of brain area with Prussian blue staining was calculated using ImageJ software (National Institutes of Health). ICH severity was expressed as the average percentage of hemosiderin surface area on 28 sections for each brain. The presence or absence of porencephaly on sections used in ICH analysis was also recorded.

### Genome scan

Genomic DNA was obtained from 192 (CAST X B6) F_2_ mice carrying the *Col4a1*^*+/Δex41*^ mutation and genotyped at the UCSF Genomic Core Facility with a commercial single nucleotide polymorphism (SNP) panel (Illumina, San Diego, CA) which contains 646 informative SNPs for (CAST X B6) F2 progeny. A genome-wide suppressor screen was performed in R (R Core Team, 2019) using the package R/qtl (Broman et al., 2003) treating the trait as binary. Genome-wide significance was established using 1000 permutation testing (Churchill and Doerge, 1994). A confidence interval for each QTL location was then calculated using Bayesian confidence sets (Sen and Churchill, 2001). The ASD score for each animal was averaged from both eyes and treated as a dichotomous variable where score of 1 was mild, and score of 1.5 - 3 was severe. A subset (49) of these 192 mice was randomly selected in a separate mapping analysis for the muscle modifier as a quantitative trait.

### Quantitative polymerase chain reaction (qPCR)

Eyes from P0 mice were dissected in PBS, immediately transferred in RNA*later* (ThermoFisher Scientific, USA) and stored at −80°C until use. Total RNA was extracted using TRIzol Reagent (ThermoFisher Scientific) according to the manufacturer’s instructions and reverse transcribed to cDNA using iScript cDNA synthesis kit (Bio-Rad, Hercules, CA). Quantitative PCR was performed on a Bio-Rad CFX96 real-time system using SsoFast Evagreen mix (Bio-Rad) and primers are listed in Table S3. Briefly, 10 ng of cDNA and 1.25 μM primers were used per reaction in a final volume of 10 μl. Each cycle consisted of denaturation at 95°C for 5s, followed by annealing and extension at 60°C for 5s for a total of 45 cycles. All experiments were run with technical duplicates and 4-6 biological replicates were used per group. The relative expression of each gene was normalized to *Hprt1* or *Gapdh* and analyzed using the 2^−ΔΔCT^ method (Livak and Schmittgen, 2001).

### Western blot analyses

Quadriceps from P10 pups were dissected and total proteins extracted using radioimmunoprecipitation assay (RIPA) buffer (VWR, Radnor, PA) supplemented with Halt Protease and Phosphatase Inhibitor Cocktail (ThermoFisher Scientific), EDTA and 2 mM phenylmethylsulfonyl fluoride. Total proteins (10 μg) were separated on 4–15% gradient SDS-PAGE gels (Bio-Rad) and transferred to polyvinylidene fluoride (PVDF) membranes (BioRad). Membranes were blocked for 1 hr at room temperature in 5% non-fat milk in TBS containing 0.1% Tween-20 (TBST), incubated overnight at 4°C in primary antibodies diluted in 2% non-fat milk in TBST. Primary antibodies and dilutions used are as follow: rabbit anti-FN1 (Abcam #Ab2413 1:10000, Abcam), rabbit anti-pFAK (1:1000, Invitrogen #44624G), rabbit anti-FAK (1:1000, Santa Cruz #SC-557), rabbit anti-ILK (1:1000, Cell Signaling Technology #3862), mouse anti-ITGB1 (1:1000, BD Biosciences #610467), and mouse anti-GAPDH (1:500000, Millipore #MAB374). Following washes in TBST, membranes were incubated for 1 hr at room temperature with species-specific horseradish peroxidase conjugated secondary antibodies (1:10000, Jackson ImmunoResearch, West Grove, PA) diluted in 2% non-fat milk in TBST. Immunoreactivity was visualized using chemiluminescence (ECL or Luminata Forte substrate, ThermoFisher Scientific) and imaged using a ChemiDoc MP Imaging System (Bio-Rad) or exposed to X-ray films. Densitometric analyses were performed on low exposure images using the Quantity One software (Bio-Rad).

### Statistics

Statistical analyses and graphs were prepared using GraphPad Prism v8.0 Software (GraphPad, La Jolla, CA). Multiple comparisons were carried using one-way ANOVA followed by Tukey post-test or Kruskal-Wallis test followed by Dunn’s post-test. *p* values less than 0.05 were considered statistically significant.

## ACKNOWLEDGEMENTS

We thank members of the Gould Laboratory for their critical reviews of the manuscript. We also thank Cassandre Labelle-Dumais for helpful discussion and guidance with the muscle studies.

## AUTHOR CONTRIBUTIONS

Conceived and designed the experiments: D.B.G and M.M. Performed experiments: M.M, T.P, M.J, and K.H. Data analysis: M.M, T.P, M.J, K.H, S.S and D.B.G. Contributed reagents, materials, and/or analysis tools: S.S. Writing of the manuscript: M.M and D.B.G.

---

**Figure S1. Subcongenic Line 1 did not confer the modifier effect.** (**A-B**) A ~10 Mb proximal portion of the CAST *ModGS1* locus (52418578-61808145bp on Chr 1, Ensembl GRCm38.p6) showed no protective effect for ASD (A) or skeletal myopathy (B) phenotypes seen in *Col4a1*^*+/Δex41*^ mice. Sample size indicated in parentheses. Data are presented as mean ± SEM. one-way ANOVA followed by Tukey’s multiple comparison test.

**Table S1. List of protein-coding and non-coding genes in the *ModGS1* locus.**

**Table S2. SNPs, small InDels and structural variants in *Fn1***.

**Table S3. Primer sequences for quantitative PCR.**

